# The effect of vitamin D and calcium supplementation on gut microbiota composition in adolescents living with HIV in Southern Africa

**DOI:** 10.1101/2025.11.20.689294

**Authors:** Matthias Hauptmann, Nyasha V Dzavakwa, Mizinga J Tembo, Molly Mazombwe-Chisenga, Christoph Leschczyk, Gajanan Thirumoolanathan, Karen Sichibalo, Tafadzwa Madanhire, Sarah L Rowland-Jones, Paul Kelly, Katharina Kranzer, Suzanne Filteau, Celia L Gregson, Victoria Simms, Lackson Kasonka, Rashida A Ferrand, Ulrich E Schaible

## Abstract

Adolescents living with HIV are at risk of delayed growth and of increased susceptibility to infections, potentially linked to vitamin D insufficiency, which is common in people living with HIV.

We analyzed rectal swab microbiota compositions before and after 48 weeks’ supplementation with high-dose vitamin D and calcium, or placebo, embedded in a randomized controlled trial conducted in Zambia and Zimbabwe (Pan African Clinical Trials Registry PACTR20200989766029). In a sub-cohort of 244 participants, we first assessed baseline associations of the rectal swab microbiota with vitamin D metabolites, parathyroid hormone (PTH), and bone mineral density (BMD). We then analyzed longitudinal changes in rectal swab microbiota and intestinal barrier integrity during supplementation and examined whether the baseline microbiota influenced the treatment response.

At baseline, anaerobic taxa within the class Clostridia were positively associated with PTH, 1,25(OH)2D, and BMD, and negatively associated with 24,25(OH)2D. After supplementation, Shannon diversity was decreased (β^Std^=-0.495, CI [-0.802, -0.188], p=0.0018), median Bray-Curtis dissimilarity from baseline samples was increased (β^Std^=0.399, CI [0.086, 0.712], p=0.013), and abundance of Clostridia_vadinBB60_group was decreased (log_2_fc=-1.715, CI [-2.596, -0.835], q=0.043) in the intervention arm. The intervention did not affect the intestinal barrier integrity, measured by intestinal fatty acid binding protein (iFABP) concentration in blood plasma. On average, PTH concentrations decreased and BMD increased during supplementation. However, higher baseline abundance of Clostridia-member UCG-009 predicted a smaller reduction in PTH concentrations during supplementation (log_2_fc^std^=1.200 CI [0.656, 1.744], q=0.012), while higher baseline abundances of *Enterococcus* (log_2_fc^std^=-1.220 CI [-1.807, -0.632], q=0.022) and *Fournierella* (log_2_fc^std^=-1.122 CI [-1.694, -0.551], q=0.025) predicted a smaller increase in BMD.

In conclusion, there was a reduction of an anaerobic taxon of the class Clostridia following vitamin D supplementation. Further, the microbiota appeared to moderate the effect of the intervention on PTH and bone density.

## Introduction

Adolescents with perinatally-acquired HIV are at risk of impaired growth and intercurrent infections, even when viral loads are suppressed by antiretroviral therapy (ART). These outcomes are influenced by multiple factors, and vitamin D insufficiency has been proposed as one possible contributor.(1) Vitamin D insufficiency is also associated with reduced intestinal barrier integrity, disturbed intestinal microflora, and intestinal inflammation, as shown in the context of inflammatory bowel disease.(2)

Pro-vitamin D3 (cholecalciferol) is primarily synthesized in the skin, when exposed to ultraviolet (UV-B) light, and is subsequently hydroxylated to 25-hydroxyvitamin D3 (25(OH)D) in the liver. Dietary intake plays a minor role in maintaining circulating 25(OH)D concentrations, unless supplemented. The circulating active form, 1,25-dihydroxyvitamin D (1,25(OH)2D), is produced mainly from 25(OH)D in the kidneys.(3) Calcium homeostasis and bone mineralization are primarily regulated by parathyroid hormone (PTH) and 1,25(OH)2D. PTH enhances bone resorption and renal calcium reabsorption, while 1,25(OH)₂D inhibits PTH synthesis and promotes intestinal calcium absorption. Together, they maintain homeostatic serum calcium concentrations through a finely regulated feedback mechanism.(4)

The gut microbiota is a complex community of microorganisms residing in the intestinal tract. It fulfils many functions within the human body, including metabolization of food, detoxification of metabolites such as bile acids, regulation of intestinal barrier integrity, resistance against colonization by pathogens, and regulation of immunological trajectories. The composition of the microbiota is variable between, but relatively stable and resilient within individuals. It is acquired in early life from family members and additionally shaped by dietary habits, disease, drug exposure, physical activities, and many other factors.(5) People living with HIV often have decreased intestinal barrier integrity, and generally less diverse microbiota, reduced in obligate anaerobes, and increased in Proteobacteria, a phylum that contains many taxa with facultative pathogenic potential.(6, 7) Antiretroviral therapy only partially restores microbiota homeostasis and intestinal barrier integrity.(8, 9)

We first analyzed cross-sectional baseline data from a randomized controlled trial of vitamin D and calcium supplementation to determine whether vitamin D and calcium metabolism regulation is associated with microbiota-patterns or intestinal barrier integrity in adolescents with perinatal HIV infection. Subsequently, we analyzed whether vitamin D and calcium supplementation for 48 weeks altered either microbiota or intestinal barrier integrity, and whether microbiota alterations influenced the skeletal response to supplementation.

## Material and Methods

### Study Design

This study was nested within the VITALITY trial, a randomized double-blind placebo-controlled trial of high-dose vitamin D with calcium supplementation in adolescents aged 11-19 years living with perinatally acquired HIV and taking ART for at least six months, aiming to improve musculoskeletal health. Participants were recruited from two public sector outpatient HIV clinics: the Women and Newborn Hospital at the University Teaching Hospital, Lusaka, Zambia and at the Children’s Hospital at Sally Mugabe Central Hospital, Harare, Zimbabwe. Randomization strategy, treatment, and the complete follow-up investigations have been described within the trial protocol.(10, 11) Briefly, participants in the intervention arm received 20,000 IU cholecalciferol (vitamin D3) weekly and 500 mg calcium carbonate daily for 48 weeks. Placebos were identical in appearance and taste to the products in the intervention arm. Participants were followed up at 2, 12, 24, and 36 weeks, for supply of trial drug and follow-up questionnaires and then, at 48 weeks for bone densitometry, anthropometric measurements, follow-up questionnaires, rectal swab- and blood collection. In the main trial, no overall differences in bone outcomes were observed; however, among participants with vitamin D insufficiency, supplementation modestly improved TBLH-BMD and LS-BMAD. No severe adverse events related to supplementation occurred.(12)

Of the 842 participants in the VITALITY trial, 248 were invited to participate in the microbiome sub-study. In Harare, these consisted of the first 121 + 3 additional recruits. In Lusaka, to coordinate with sampling for other VITALITY sub-studies, two batches of 61 and 63 consecutive VITALITY recruits were invited.

### Sample collection

Rectal swab samples were collected at baseline and 48 weeks by study nurses using flocked swabs. As part of quality control, only samples visibly stained with fecal matter were used. Samples were then placed in DNA/RNA Shield buffer (swabs and buffer from ZymoResearch, Irvine, US) and stored at -20 °C.

At baseline and 48 weeks, EDTA-plasma was collected at both study sites. In Lusaka, to coordinate with other VITALITY sub-studies, heparin-plasma was collected during peripheral blood mononuclear cell (PBMC) isolation (results of PBMC-analyses will be reported separately).

### Questionnaire data

Socio-economic status (SES) was derived from variables measuring household asset ownership and income, using factor analysis. Dietary intake of vitamin D and calcium were assessed based on the FANTA food frequency tool which is applicable to sub-Saharan Africa.(13) Respiratory symptoms were defined as any report of fever, cough, nasal congestion, sore throat, sneezing, muscle aches, headache, loss of smell/taste, difficulty breathing, rapid heartbeat, or chest pain at all visits. Cotrimoxazole treatment was reported as indicated in baseline questionnaires; other antibiotic intake was reported as at all visits.

### Blood measurements

In Harare, a finger-prick sample was collected for CD4 count testing and was analyzed using an Abbott CD4 PIMA analyzer. In Lusaka, CD4 count was measured in EDTA blood using the Aquois CL flow cytometer (Beckman Coulter) or an Abbott CD4 PIMA analyzer. HIV-1 viral load testing was done using the Qiagen rotor gene Q, Hologic Panther or GeneXpert in Lusaka and the Roche COBAS Ampliprep/COBAS Taqman48 in Harare. EDTA blood plasma aliquots were stored at -80 °C and shipped to the University of East Anglia for mass spectrometry testing of 25(OH)D, 1,25(OH)D, 24,25(OH)D, and PTH.(14)

Aliquots of EDTA blood plasma (Harare samples) or heparin blood plasma (Lusaka samples) were frozen at -80 °C and transported to Germany on dry ice. Prior to analysis, samples were rapidly thawed (30 min at 37 °C in a water bath) and supplemented with additional 4.7 mM sodium EDTA to avoid precipitation. Intestinal fatty acid binding protein (iFABP) concentrations were measured using Duo Set iFABP ELISA-Kits (R&D Systems, Minneapolis, US) using sterile 50 mM Tris, 10 mM CaCl2, 150 mM NaCl supplemented with 20% goat serum and 1% Triton X 100 as sample buffer, and otherwise following the manufacturer’s protocol. iFABP is a cytosolic protein of enterocytes, and its release into plasma reflects epithelial injury and impaired intestinal barrier integrity. (15)

### Bone density measurements

Total body less-head bone mineral density (TBLH-BMD) and lumbar spine-bone mineral apparent density (LS-BMAD) were measured using Dual-energy X-ray absorptiometry (DXA) scanning. The GE-Lunar iDXA and the Hologic Discovery DXA systems were used in Harare and Lusaka, respectively. The European spine phantom (a semi-anthropomorphic calibration standard (16) was scanned for cross-calibration of both machines every 6 months, in addition to daily quality control scanning of the manufacturer provided phantom on each machine.

### 16S rRNA gene sequencing

DNA extraction was performed in batches of 18 samples, including 3 participants per trial arm with samples from 3 time points per participant (baseline, 48 weeks, and 96 weeks. Data from the 96 weeks follow-up will be reported separately). A water control and a microbial mock control sample (ZymoResearch, Irvine, US) were included in each extraction batch. DNA was extracted using a commercial kit (ZymoBIOMICS DNA Miniprep Kit, ZymoResearch), including a bead beating step with mixed 0.1 mm and 0.5 mm beads (ZymoResearch) on a FastPrep device (MP Biomedicals, Irvine, US) at maximum speed (5 times 50 sec beating + 5 min cooling). DNA was eluted in water and stored at -80 °C.

The 16S rRNA V3-V4 region was amplified by 25 PCR cycles using standard 341-fw (CCTACGGGNGGCWGCAG) and 805-rv (GACTACHVGGGTATCTAATCC) primers (17), elongated by Illumina adapter sequences (Eurogentec, Seraing, Belgium). Amplification was performed with maximal 10 ng of DNA template using High Fidelity Hot Start Polymerase (Kapa Biosystems, Wilmington, US), followed by DNA-clean-up using magnetic beads (MagnaMedics, Aachen, Germany). Index-PCR was performed using 8 PCR cycles with primers containing barcode indexes (Illumina DNA/RNA UDI Tag 96 Idx, Illumina, San Diego, US) and using High Fidelity Hot Start Polymerase, followed by DNA-clean-up using magnetic beads. DNA concentrations were quantified using picogreen assays (Thermofisher, Waltham, US). Samples were diluted to equal DNA concentrations and pooled as described in Illumina’s 16S rRNA sequencing protocol. Pooled libraries were sequenced on a NextSeq 2000 instrument using P1 600 cycles reagent kits and a target of 40% phiX spike-in.

Sequences from each sequencing run were demultiplexed, paired reads were merged, trimmed from primer sequences, truncated (fw reads at 290 bp, rv reads at 240 bp), and quality filtered (max. expected errors=2). The resulting sequences were denoised and chimera-filtered to obtain amplicon sequence variants using QIIME2 with DADA2 version 2021.8.0. (18, 19) After denoising, feature-tables and representative sequences from all sequencing runs were combined using the QIIME2 commands ‘feature-table merge’ and ‘feature-table merge-seqs’. A feature-classifier was trained with the QIIME2 command ‘feature-classifier fit-classifier-naïve-bayes’ with the SILVA database version 138.99 (20) and used to classify the merged representative sequences by taxonomy.

### Data analysis

Vitamin D insufficiency was defined as 25(OH)D plasma concentrations <75 nmol/L, based on an analysis we published previously.(21) Dry season was defined as May to October and rainy season as November to April.

TBLH-BMD Z-scores and LS-BMAD Z-scores were determined in relation to age- and sex-matched European reference data, following ISCD recommendations.(22)

Data were analyzed using R Studio version 2024.12.0 with R version 4.4.1. Relative risks were calculated using the epitools package (version 0.5-10.1). For 16S rRNA sequencing data, phyloseq version 1.48.0 was used. Shannon diversities were calculated using the vegan package (version 2.6-8). The Bray-Curtis value of a sample is the median of the Bray-Curtis dissimilarities to all baseline samples. Scaling to Z-scores and linear regression models were done using R base package. Linear mixed-effects models were built using function lmer of lme4 package version 1.1-35.5. To analyze whether intervention influenced the microbiota α- or β-diversity or the intestinal barrier integrity, we used linear mixed models to fit Z-score transformed Shannon index, median Bray-Curtis dissimilarity to baseline samples, or iFABP concentrations in blood plasma by time point × arm interaction, while time point, arm, and study site were kept as fixed effects and participant as random effects covariates. Linear Models for Differential Abundance Analysis (LinDA) was used from the MicrobiomeStat package (version 1.2) to determine differential taxa abundances.(23) To analyze the intervention effect on the composition of the rectal swab microbiota, we used LinDA to model taxa abundances by time point × arm interaction, using time point, arm, and study site as fixed effects and participant as random effects covariates. To evaluate whether the microbiota influenced treatment response, we calculated Z-scores for PTH, 25(OH)D, and 1,25(OH)2D based on the internal distribution of values at baseline and 48 weeks. Bone outcomes (TBLH-BMD and LS-BMAD) were already available as age- and sex-specific Z-scores based on external European reference populations. To enable direct comparison of beta-coefficients across all variables, these externally-referenced bone Z-scores were further standardized to internal Z-scores, placing them on the same scale as the metabolite Z-scores. The differences between 48 week and baseline Z-scores (ΔZ) were then calculated for each variable and used as indicators of the strength of the treatment response. LinDA was used to determine differential taxa abundances at baseline by ΔZ values in bivariate regression models, using study site as a covariate. False discovery rate (FDR-) adjustment was used for multiple comparison corrections. Permutational multivariate analysis of variance (PERMANOVA) was conducted using the adonis2 function from the vegan package (version 2.6-8) on a Bray-Curtis dissimilarity matrix. To account for the longitudinal structure of the data, permutations were carried out within each participant rather than across participants, preserving the paired nature of the repeated measures. For models that included Z-scored variables, results were reported as standardized β-coefficients (β^std^) for linear and linear mixed regression models, or as standardized log2 fold-change values (log_2_fc^std^) for LinDA models. 95% confidence intervals of all models were calculated using base R. Heatmaps were made using pheatmap (version 1.0.12).

### Ethical considerations

Written informed consent to participate in the parent VITALITY trial was obtained in the local language from participants’ guardians with written assent from participants aged <18 years. Participants aged ≥18 years provided independent consent. Ethical approval was granted by the Biomedical Research and Training Institute Institutional Review Board (AP158/2020), the Harare Central Hospital Ethics Committee (HCHEC 030320/12), the Medical Research Council of Zimbabwe (A/2626), the University of Zambia Biomedical Research Ethics Committee (1116–2020), and the London School of Hygiene and Tropical Medicine Ethics Committee (22030). The VITALITY trial is registered with the Pan African Clinical Trials Registry PACTR20200989766029.

## Results

### Study overview

From 842 adolescents within the VITALITY main trial, 248 (29%) were invited to participate in the microbiome sub-study. Of those, two refused participation and the rectal swab samples of two others did not meet the quality criteria. Thus 244 participants were enrolled and provided rectal swabs at baseline. Of those, 120 were randomly allocated to the intervention arm and 124 into the control arm. Subsequently, 109 participants in the intervention arm and 115 participants in the control arm had a 48 weeks study visit (Fig. 1). Participants in Harare were enrolled between 2021-02-04 and 2021-05-18, those in Zambia between 2021-07-12 and 2021-07-29 or between 2021-08-23 and 2021-09-22. Age, sex, BMI, 25(OH)D concentrations and HIV viral load at baseline were similar in participants assigned to intervention and control. While in Lusaka no participant received cotrimoxazole, around half of the participants in Harare received it continuously as a prophylactic therapy. Due to the staggered recruitment timeline, all participants in Lusaka had the baseline visit within the dry season, while this was the case for only two participants in Harare (both in intervention arm, one without rectal swab at 48 weeks), all other participants in Harare had their baseline visit during the rainy season. Few participants received antibiotics other than cotrimoxazole during the 48 weeks of intervention, and numbers were comparable between the groups. More participants in the intervention compared to control arm reported any respiratory symptoms during the 48 weeks of intervention (21.7% vs 12.1%, p=0.046; Table 1).

**Figure 1:**
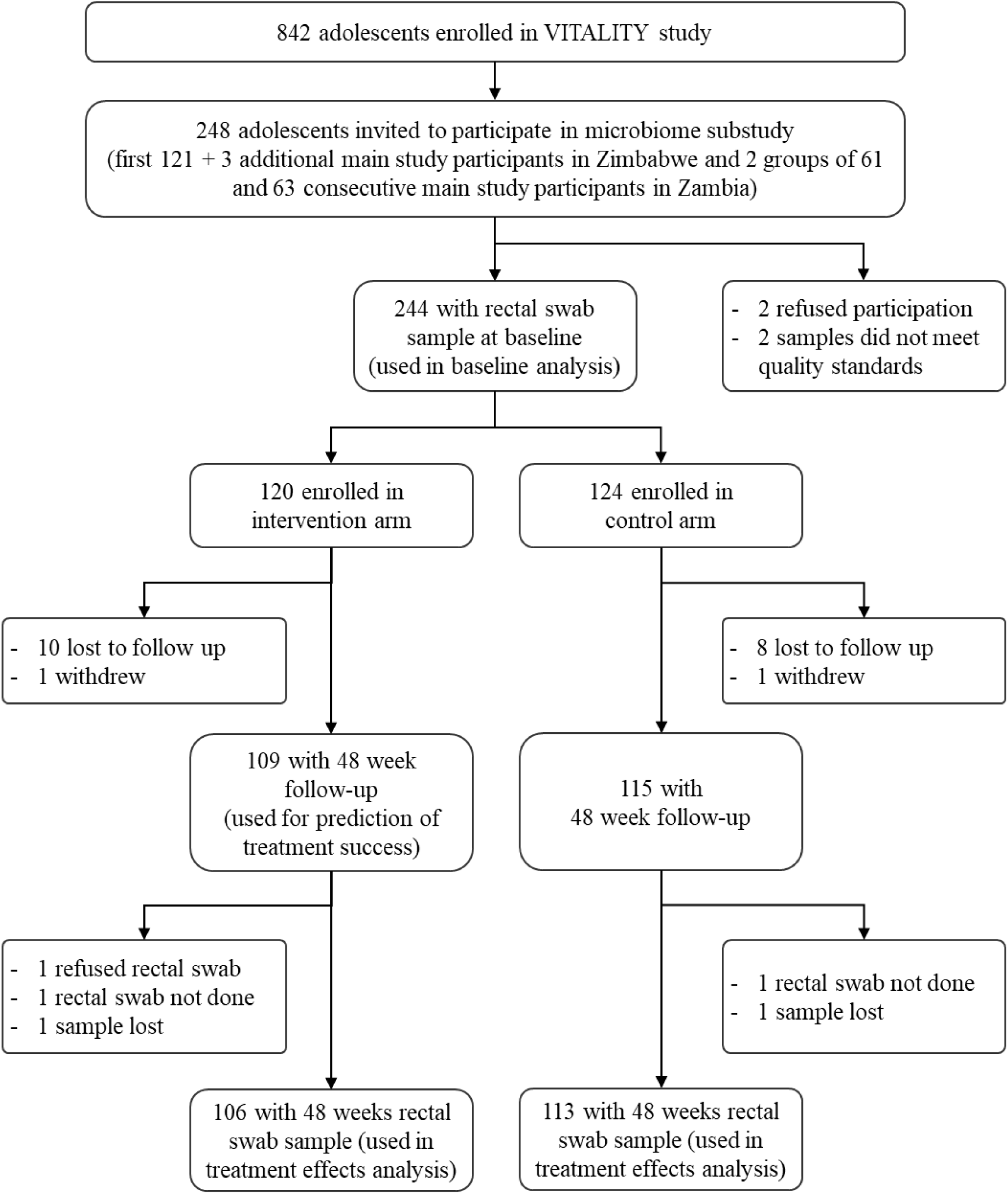
Trial flow chart.

**Table 1.** Demographic and clinical data.

| Participant characteristic | Total (244 participants) | Intervention arm (120 participants) | Placebo arm (124 participants) |
| --- | --- | --- | --- |
| female sex, n (%) | 119 (48.8%) | 59 (49.2%) | 60 (48.4%) |
| age, years (baseline), median (IQR) | 15.5 [13.5, 18.0] | 15.5 [13.5, 17.5] | 16.0 [13.0, 18.0] |
| BMI<sup>*1</sup>, kg/m<sup>2</sup> (baseline), median (IQR) | 18.4 [16.4, 20.2] | 18.4 [16.4, 20.3] | 18.3 [16.4, 20.2] |
| 25(OH)D<sup>*1</sup> nmol/L (baseline), median (IQR) | 67.3 [56.5, 81.9] | 66.9 [54.9, 79.9] | 68.8 [58.1, 84.5] |
| viral load copies/ml (baseline), median (IQR) | 20.0 [0.00, 60.5] | 20.0 [0.0, 65.2] | 20.0 [0.0, 59.0] |
| any antibiotic use except cotrimoxazole (0-48 weeks), n (%) | 10 (4.1%) | 6 (5.00%) | 4 (3.23%) |
| any respiratory symptoms (0-48 weeks), n (%) <sup>*2</sup> | 41 (16.8%) | 26 (21.7%) | 15 (12.1%) |
| country Zimbabwe, n (%) | 121 (49.6%) | 60 (50%) | 61 (49.2%) |
| cotrimoxazole (baseline), n (%) | 62 (25.4%) | 32 (26.7%) | 30 (24.2%) |
| in Lusaka, n (%) | 0 (0%) | 0 (0%) | 0 (0%) |
| in Harare, n (%) | 62 (51.2%) | 32 (53.3%) | 30 (49.2%) |
| baseline sample during dry season n (%) | 125 (51.2%) | 62 (51.7%) | 63 (50.8%) |
| in Lusaka, n (%) | 123 (100%) | 60 (100%) | 63 (100%) |
| in Harare, n (%) | 2 (1.65%) | 2 (3.33%) | 0 (0%) |

<sup>\*1</sup> Abbreviations: BMI, body mass index; 25(OH)D, 25-hydroxyvitamin D
<sup>\*2</sup> intervention vs. placebo arm respiratory symptoms. chi.square: p=0.046

### Quality of 16S rRNA gene sequencing results

Eight of 463 DNA samples from rectal swabs produced less than 50,000 high-quality reads (i.e. after denoising), and the microbial mock community sample included in one extraction batch disagreed with the theoretical composition. Sequencing of samples from the affected batch and of samples with less than 50,000 reads was repeated. Afterwards, the minimum read-number after denoising was 52,355 (median 148,984; IQR: 108,983 - 205,411), and the microbial mock communities from all batches matched the expected theoretical composition (Fig. S1).

### Baseline microbiota and barrier integrity: description and associations

*Firmicutes* was the most abundant phylum in rectal swabs at baseline, followed by *Bacteroidota* and *Actinobacteria*. *Prevotella* was the most abundant genus, followed by *Faecalibacterium*, *Peptoniphilus*, and *Bacteroides* (Fig. 2). We observed significant differences in microbiota compositions between the two study sites at genus level (R^2^=0.021, p<0.001 by permutational multivariate analysis of variance, PERMANOVA), while no difference was observed at phylum level (R^2^=0.0045, p=0.346 by PERMANOVA). There were no differences in microbiota compositions at genus level between participants from Harare, who received or did not receive prophylactic cotrimoxazole treatment (R^2^=0.0074, p=0.52 by PERMANOVA).

**Figure 2:**
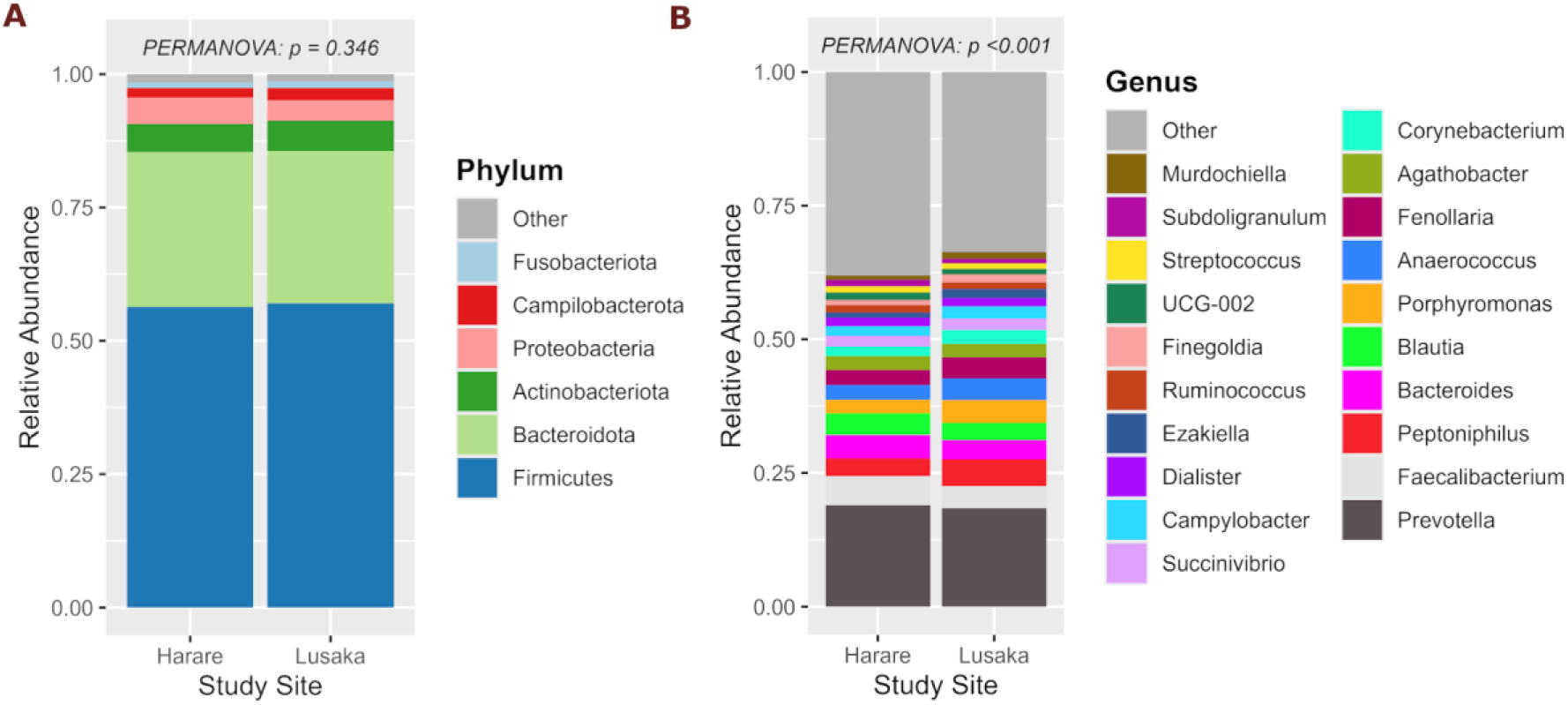
Microbiota compositions between study sites. Mean relative abundances of the 6 most abundant phyla **(A)** or the 20 most abundant genera **(B)** within rectal swab microbiota are shown, stratified by study site. p-values indicate results of permutational multivariate analysis of variance (PERMANOVA) comparing community compositions across study sites among all taxa at - or genus level, respectively. **Sample size:** Harare, n=121; Lusaka, n=123.

We analyzed differential bacterial taxa abundances in the rectal swab microbiota at baseline using Linear Models for Differential Abundance Analysis (LinDA). We conducted bivariate analyses (i.e., one analysis per independent variable using study site as a covariate). The prescribed antiretroviral drugs were not associated with differential abundances within the 6 most abundant phyla or the 20 most abundant genera within the rectal swab microbiota. However, using linear regression models adjusted for study site, we found that Tenofovir disoproxil fumarate (TDF) was associated with higher intestinal fatty acid binding protein (iFABP) (β^std^=0.607 CI [0.303, 0.911], q=0.0036), while Abacavir (ABC) was associated with lower iFABP concentrations (β^std^=-0.905 CI [-1.416, -0.395], q=0.0091) in blood plasma, indicating lower or greater intestinal barrier integrity, respectively. Lopinavir/ Ritonavir (LPV.r) was associated with lower Shannon diversity (β^std^=-0.888 CI [-1.459, -0.318], q=0.025) (Suppl. Fig. 2). We found lower abundance of the phylum *Fusobacteriota* (log_2_fc=-2.893 CI [-4.049, -1.738], q=8.62e-5) and higher abundance of the genus *Anaerococcus* (log_2_fc=1.039 CI [0.525, 1.552], q=0.016) in males compared with females. We also found increased abundances of the phylum *Actinobacteriota* (log_2_fc^std^=0.567 CI [0.357, 0.778], q=2.94e-5) as well as the genus *Corynebacterium* (log_2_fc^std^=1.218 CI [0.804, 1.631], q=8.11e-6) as age increased. BMI was negatively associated with the abundance of the genus *Anaerococcus* (log_2_fc^std^=-0.497 CI [-0.756, -0.237], q=0.025). Abundance of the genus *Porphyromonas* was negatively associated with blood plasma concentrations of 25(OH)D (log_2_fc^std^=-0.668 CI [-1.030, -0.306], q=0.028), and the genus *Subdoligranulum* was positively associated with plasma concentrations of active vitamin D (1,25(OH)2D) (log_2_fc^std^=0.437 CI [0.174, 0.699], q=0.043) and PTH (log_2_fc^std^=0.453 CI [0.191, 0.716], q=0.038), but negatively associated with concentrations of the vitamin D degradation product 24,25(OH)2D (log_2_fc^std^=-0.523 CI [-0.827, -0.220], q=0.038). Total body less head bone mineral density Z-score (TBLH-BMD Z-score) was positively associated with abundance of Unclassified Clostridiales Group 002 (UCG-002), a genus within the family *Oscillospiraceae* (log_2_fc^std^=0.501 CI [0.209, 0.793], q=0.038), and negatively associated with the genera *Streptococcus* (log_2_fc^std^=-0.572 CI [-0.910, -0.234], q=0.040) and *Anaerococcus* (log_2_fc^std^=-0.483 CI [-0.746, -0.219], q=0.028). Lumbar spine bone mineral apparent density Z-score (LS-BMAD Z-score) showed associations consistent, but weaker, than TBLH-BMD Z-score with the three aforementioned genera. Shannon diversity was positively associated with TBLH-BMD Z-score (β^std^=-0.243 CI [0.138, 0.348], q=0.00047). No FDR-adjusted significant associations were seen between any of the analyzed baseline demographic, anthropometric, clinical, dietary, biochemical, or DXA-derived bone variables with blood plasma concentrations of iFABP, a marker for intestinal barrier integrity, or Bray-Curtis dissimilarity (Figure 3).

**Figure 3:**
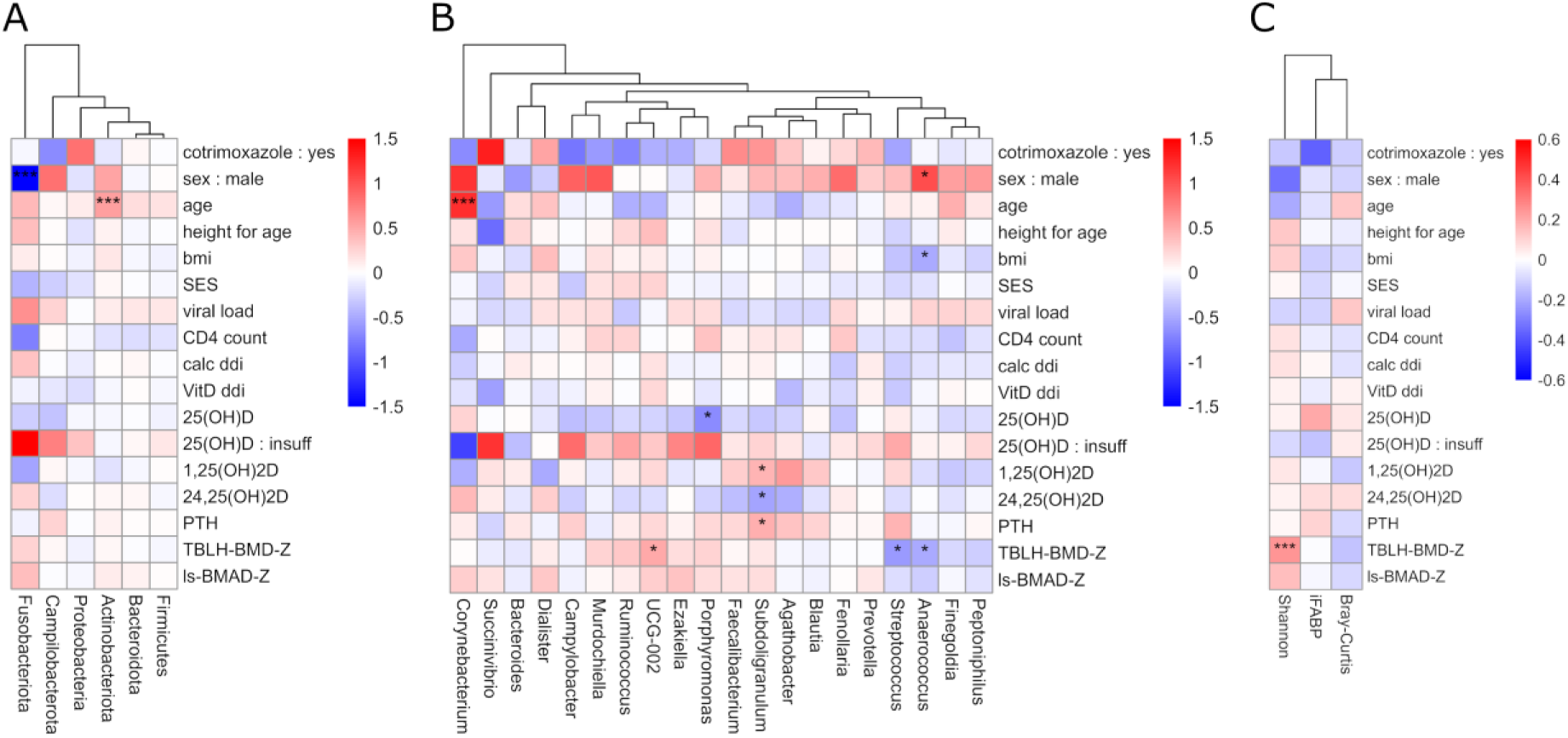
Baseline microbiota associations with clinical data. **A, B)** Differential abundances in baseline microbiota by the indicated variables were assessed using LinDA, adjusting for study site. Results are shown for the 6 most abundant phyla **(A)** and the 20 most abundant genera **(B)**. Colors represent log₂ fold changes. **C)** Associations of Shannon diversity, Bray-Curtis dissimilarity, and plasma iFABP concentrations with the indicated variables were assessed using linear regression, adjusting for study site. Colors represent β-coefficients. **A-C)** Continuous variables were standardized prior to modeling. **Sample size**: n=244. Asterisks denote FDR-adjusted significance: * q<0.05, *** q<0.001. **Abbreviations**: LinDA, Linear Models for Differential Abundance Analysis; SES, socioeconomic status; ddi, daily dietary intake; PTH, parathyroid hormone; TBLH-BMD-Z, total body less head bone mineral density Z-score; LS-BMAD-Z, lumbar spine bone mineral apparent density Z-score; iFABP, intestinal fatty acid binding protein.

### Impact of vitamin D + calcium intervention on taxa abundances, α- and β-diversity and intestinal barrier integrity

We tested the effects of high dose vitamin D + calcium treatment on the composition of the rectal swab microbiota using LinDA. Treatment effects indicate a time point by arm interaction from models including time point, arm, and study site as fixed, and participant as random effects. Since we expected stronger effects of vitamin D supplementation in participants with vitamin D insufficiency, we applied a subgroup analysis including only participants with vitamin D insufficiency at baseline (n=143 participants), in addition to the analysis in the entire cohort. After FDR-adjustment, we did not find significant associations within the 6 most abundant phyla or the 20 most abundant genera in either the entire cohort or the subgroup. Trends indicated decreased abundances of the genera *Subdoligranulum* (log_2_fc=-0.951 CI [-1.614; -0.288], q=0.107 in entire cohort) and UCG-002 (log_2_fc=-0.880 CI [-1.557; -0.203], q=0.115 in the entire cohort), as well as increased abundance of *Finegoldia* (log_2_fc=-1.065 CI [0.248; 1.881], q=0.220 in the 25(OH)D-insufficient subgroup) in the intervention arm (Suppl. Figure 3). When the analysis was extended to all genera with a prevalence >0.1, Clostridia_vadinBB60_group, an unclassified genus within the class Clostridia, was significantly reduced upon intervention in the entire cohort (log_2_fc=-1.715 CI [-2.596, -0.835], q=0.043), while the reduction did not reach significance in the 25(OH)D-insufficient subgroup (log_2_fc=-1.897 CI [-2.993, - 0.801], q=0.105). We further analyzed whether the intervention influenced the microbiota α- or β-diversity or the intestinal barrier integrity. We found a significant decrease in Shannon diversity (β^Std^=-0.495, CI [-0.802, -0.188], p=0.0018 in the entire cohort; β^Std^=-0.574, CI [-0.958, -0.189], p=0.0040 in the 25(OH)D-insufficient subgroup). We found increased median Bray-Curtis dissimilarity by treatment (β^Std^=0.399, CI [0.086, 0.712], p=0.013 in the entire cohort; β^Std^=0.417, CI [0.008, 0.826], p=0.048 in the 25(OH)D-insufficient subgroup) using the linear mixed model approach, but no significant effect on microbiota composition using PERMANOVA; (R^2^=0.0025, p=0.071 in the entire cohort; R^2^=0.0028, p=0.284 in the 25(OH)D-insufficient subgroup). We found no evidence of a treatment effect on iFABP concentrations, within either the entire cohort or the 25(OH)D-insufficient subgroup. (Figure 4)

**Figure 4:**
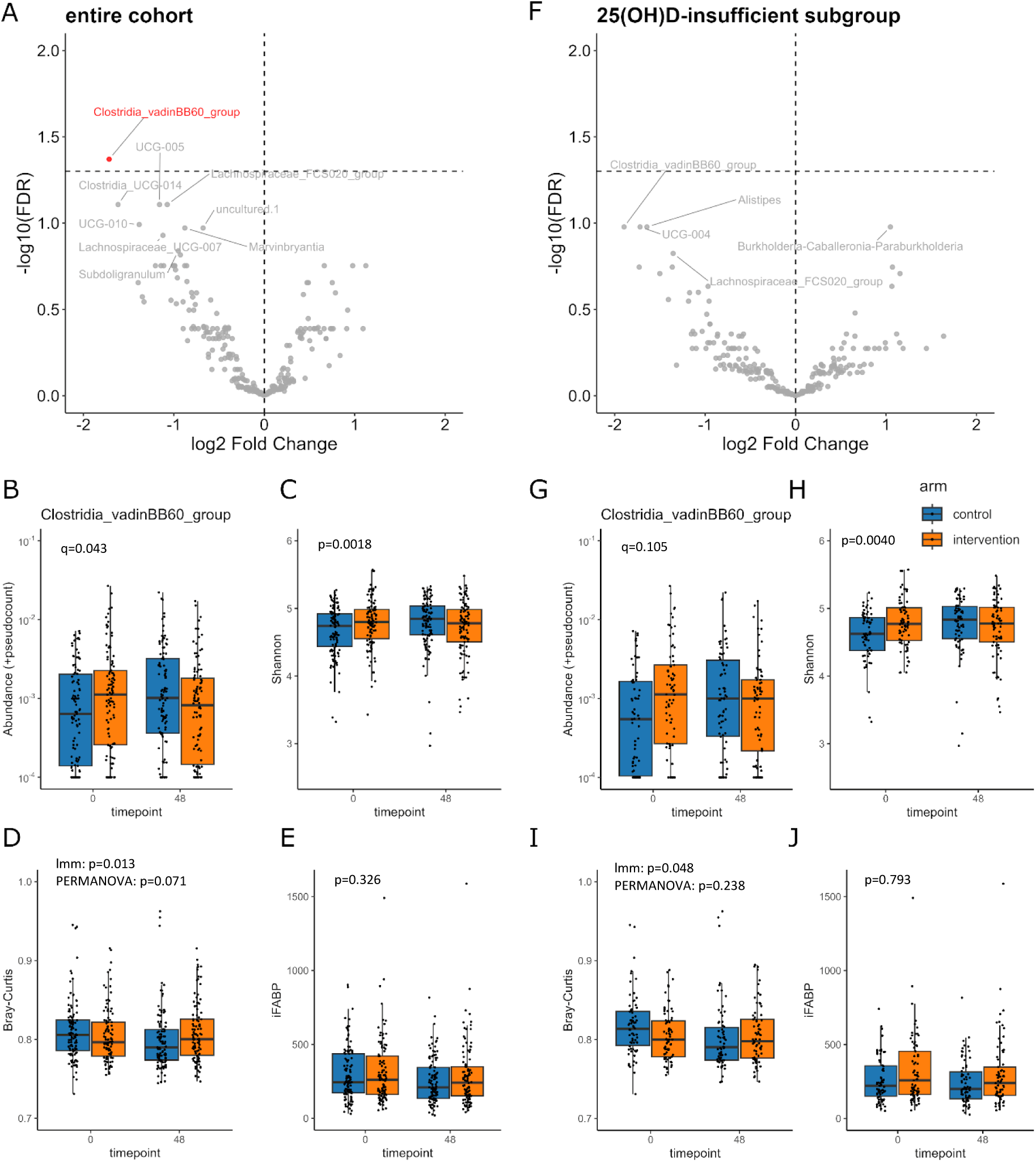
Effects of vitamin D + calcium supplementation on microbiota compositions. Analyses include all participants with baseline and 48-week samples **(A-E)** or participants with 25(OH)D-insufficiency at baseline **(F-J)**. **A, F)** Differential genus-level abundances in rectal swab microbiota, associated with treatment, were assessed using LinDA, based on treatment by arm interaction^*1^. Volcano plots display log₂ fold changes and q-values. Genera with q<0.15 are labeled; red label indicates genus with q<0.05. The horizontal lines indicate the FDR threshold (q=0.05). The vertical line indicates no change (log₂ fold change=0). **B, G)** Abundance of Clostridia_vadinBB60_group across time points and treatment arms; q-values correspond to results given in A and F. **C, H)** Shannon diversity index across time points and treatment arms. p-values indicate results of linear mixed models for treatment effect, based on treatment by arm interaction^*1^. **D, I)** Median Bray-Curtis dissimilarity to baseline across time points and treatment arms. p-values indicate results of linear mixed models as described for C and H or using PERMANOVA^*2^ **E, J)** Plasma iFABP concentrations across time points and treatment arms. p-values indicate results of linear mixed models as described for C and H. **Sample sizes**: A–E: n=438 samples (219 participants); F–J: n=286 samples (143 participants) **Abbreviations**: LinDA, Linear Models for Differential Abundance Analysis; PERMANOVA, permutational multivariate analysis of variance; iFABP, intestinal fatty acid binding protein. ^*1^ Linear models were adjusted for time point, arm, and study site as fixed effects and participant as a random effect to account for repeated measures. ^*2^ PERMANOVA was adjusted for time point, arm, and study site as fixed effects, with permutations restricted within participants.

### Identification of taxa that moderate the treatment response

We were interested in whether the microbiota could affect the strength of the treatment response. Therefore, we selected five parameters as indicators of treatment response: 25(OH)D and 1,25(OH)2D, to test whether the microbiota influences intestinal uptake of vitamin D or 1α-hydroxylation of 25(OH)D resulting in active vitamin D, PTH, as an important regulator of bone accrual and serum calcium concentrations, and TBLH-BMD Z-score as well as LS-BMAD Z-score, as they were the primary and secondary outcomes of the VITALITY trial. These variables were Z-transformed and changes between 48-weeks and baseline samples (ΔZ) were used as proxies for treatment response. We conducted differential abundance analyses on phylum- and genus level to explore whether baseline abundances of microbiota members were associated with treatment response. After FDR-adjustment, we did not find any significant associations of ΔZ-values with any of the 6 most abundant phyla or the 20 most abundant genera, in either the entire cohort or in those with 25(OH)D insufficiency (Figure S4). We extended the analysis to all genera with prevalence >0.1 and found significantly lower LS-BMAD-ΔZ associated with abundances of the genera *Enterococcus* (log_2_fc^std^=-1.220 CI [-1.807, -0.632], q=0.022) and *Fournierella* (log_2_fc^std^=-1.122 CI [-1.694, -0.551], q=0.025) in the entire cohort, as well as significantly higher PTH-ΔZ associated with the abundance of Unclassified Clostridiales Group 009 (UCG-009), a genus within the family *Butyricicoccaceae* (log_2_fc^std^=1.200 CI [0.656, 1.744], q=0.012), in the 25(OH)D-insufficient subgroup (Figure 5). Similar trends were seen for the genera *Enterococcus* (log_2_fc^std^=-1.195 CI [-1.913, -0.477], q=0.204) and *Fournierella* (log_2_fc^std^=-0.970 CI [-1.660, -0.280], q=0.446) in the 25(OH)D-insufficient subgroup, and for UCG-009 in the entire cohort (log_2_fc^std^=0.796 CI [0.322, 1.270], q=0.146).

**Figure 5:**
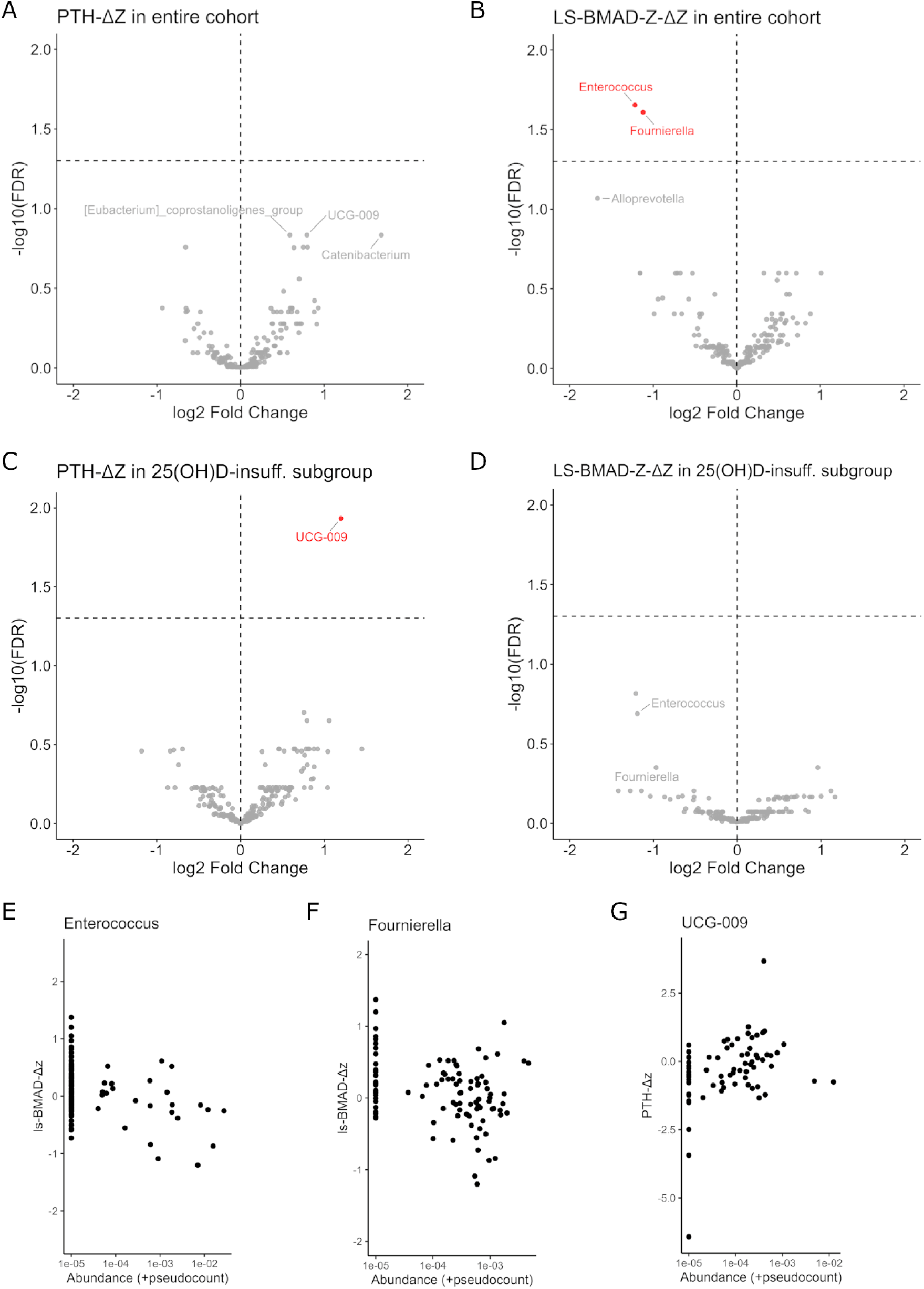
Predictors of treatment response to vitamin D + calcium supplementation. Analyses include all participants in the intervention arm **(A, B, E, F)** or those with 25(OH)D-insufficiency at baseline **(C, D, G)**. Variables 25(OH)D, 1,25(OH)2D, PTH, TBLH-BMD Z-score, and LS-BMAD Z-score were Z-transformed and changes in Z-scores between 48-weeks and baseline samples (ΔZ) were used as proxies for treatment response. Associations between genus-level abundance at baseline and ΔZ of the indicated variables were assessed using LinDA, adjusted for study site. **A–D)** Volcano plots display log₂ fold changes and q-values. Genera with q<0.15 are labeled; red labels indicate genera with q<0.05. Additionally, significant genera in the entire cohort were also labeled in the subgroup analysis. The horizontal lines indicate the FDR threshold (q=0.05). The vertical line indicates no change (log₂ fold change=0). After FDR-adjustment, no significant associations were observed for variables 25(OH)D-ΔZ, 1,25(OH)2D-ΔZ, or TBLH-BMD-ΔZ (not shown). **E-G)** Plots illustrate associations identified in A–D. **Sample sizes**: A, B, E, F: n=107-109 and C, D, G: n=73-74 participants due to incomplete TBLH-BMD Z-score and LS-BMAD Z-score data. **Abbreviations**: LinDA, Linear Models for Differential Abundance Analysis; PTH, parathyroid hormone; TBLH-BMD-Z, total body less head bone mineral density Z-score; LS-BMAD-Z, lumbar spine bone mineral apparent density Z-score.

## Discussion

We investigated baseline associations of bone density as well as concentrations of vitamin D metabolites and regulators of calcium homeostasis with microbiota composition in rectal swabs, as well as intestinal barrier integrity, in adolescents living with HIV. We then analyzed whether an intervention by vitamin D + calcium supplementation changed the microbiota or the intestinal barrier integrity, and whether the microbiota influenced the response to vitamin D + calcium supplementation.

We found associations of *Porphyromonas*, *Subdoligranulum*, Unclassified Clostridiales Group 002 (UCG-002; family Oscillospiraceae*), Streptococcus*, and *Anaerococcus*, as well as microbiota α*-*diversity with different readouts implicated in vitamin D metabolism, calcium homeostasis, and bone density at baseline. Additionally, high-dose vitamin D + calcium supplementation led to a 3.3-fold reduction of the abundance of genus Clostridia_vadinBB60_group, decreased Shannon diversity, and increased Bray-Curtis dissimilarity compared to baseline compositions. We further found that abundance of the genus UCG-009 (family *Butyricicoccaceae*) was positively associated with treatment response, as measured by PTH plasma concentrations at 48 weeks, and that abundances of the genera *Enterococcus* and *Fournierella* were negatively associated with treatment response, as measured by LS-BMAD Z-score.

*Anaerococcus* is not a typical genus within the gut, but is regularly found in the male urogenital tract, where it colocalizes with other Gram positive anaerobic cocci including *Finegoldia* and *Peptoniphilus*.(24) This aligns with the close clustering of the three genera seen in the baseline association matrix and the association of *Anaerococcus* with male sex at baseline. Concurrently, *Anaerococcus* was negatively associated with TBLH-BMD Z-score, but as TBLH-BMD Z-scores (relative to age- and sex-matched reference data) were higher in females than males (data not shown), this association was likely confounded by sex.

The genus *Subdoligranulum* and its close relative UCG-002 were positively associated with 1,25(OH)2D and PTH, or TBLH-BMD Z-score, respectively. *Subdoligranulum* is a butyrate producer and generally associated with a non-inflammatory environment, metabolic health, and intact intestinal barrier integrity.(25, 26) Further, higher Shannon diversity, which indicates a homeostatic microbiota composition, was positively associated with TBLH-BMD Z-scores, while *Porphyromonas* and *Streptococcus*, which are typically associated with dysbiosis, inflammation, and reduced obligate anaerobes, were negatively associated with 25(OH)D plasma concentrations or TBLH-BMD Z-scores, respectively. These findings indicate that homeostatic microbiota, especially their obligate anaerobic members, play a role in calcium regulation and support bone mineral density.

Vitamin D supplementation has previously been linked to dampening of pro-inflammatory immune signatures, increased intestinal barrier integrity, decreased abundances of taxa with pro-inflammatory properties, and increased abundance of the phylum Bacteroidetes, which contains many obligate anaerobes.(27, 28) There were no indications of increased barrier integrity in participants receiving vitamin D + calcium supplementation within this study. An unclassified genus within the class Clostridia (Clostridia_vadinBB60_group) was significantly reduced by 3.3-fold within the entire cohort, with a non-significant 3.7-fold decrease observed within the 25(OH)D-insufficient subgroup after 48 weeks supplementation of high-dose vitamin D3 + calcium. By trend, the closely related genera *Subdoligranulum* and UCG-002 were also reduced in the intervention group. These taxa share an anaerobic metabolism, the production of short chain fatty acids (SCFAs), and are mostly linked to anti-inflammatory conditions. The reduction of obligate anaerobes during intervention occurred alongside an increased susceptibility to respiratory tract infections, potentially due to the loss of SCFA-mediated immunomodulatory effects.(29) The reasons for the decreased abundances of the various Clostridia members are unclear and appear to contrast with previous reports that associate vitamin D supplementation with anti-inflammatory actions (27, 30, 31) and increased abundance of *Subdoligranulum.*(32) Results from other vitamin D supplementation studies reported both increased (33) and decreased (28, 34) abundances of Clostridia members, indicating a fine-tuned modulation of their metabolic capacities.

Abundance of UCG-009, another genus within the class Clostridia, predicted a smaller reduction in PTH-ΔZ during vitamin D + calcium treatment, indicating a lower PTH response in participants with higher UCG-009 abundance. Interestingly, a positive correlation of UCG-009 with bone health has been shown in a study of mice fed a High-Fructose Corn Syrup (35). While the mechanisms linking UCG-009 to bone, calcium, or vitamin D metabolism remain unclear, this study supports the broader involvement of UCG-009 in regulatory networks relevant to bone health. *Fournierella* (also member of the class Clostridia) was negatively associated with LS-BMAD Z-score informed treatment response (LS-BMAD-ΔZ) in this study, while it has been linked with increased bone health in chicken (36). *Enterococcus* potentially indicates a shift of the microbiota where abundance of anaerobic taxa is reduced. The negative association of *Enterococcus* with change in LS-BMAD Z-Score therefore supports a role of anaerobes in the response to vitamin D supplementation.

In this study, we report associations of anaerobic bacterial genera of the class Clostridia with vitamin D metabolites, regulators of calcium homeostasis, and bone mineral density using baseline associations, associations with cholecalciferol + calcium intervention vs. control, and using predictive associations with treatment response. While others have seen associations with the same or closely related taxa, the direction of the associations differs between studies.(28, 35–37) Species- or strain-specific differences between the identified taxa or differences in the microbial environments with their complex interaction networks could potentially explain this discrepancy. SCFAs have been identified as modulators of bone health via different mechanisms. First, inflammatory mediators are known to stimulate osteoclast activity (38, 39), whereas SCFAs exhibit primarily anti-inflammatory properties via reduction of inflammatory cytokine expression and induction of regulatory T cells. (40) Further SCFAs, especially butyrate, have been shown to directly inhibit osteoclast differentiation and stimulate osteoblast activity. (36, 41, 42)

We hypothesize several non-exclusive mechanisms linking abundance of obligate anaerobes and bacterial alpha-diversity with the regulatory network of vitamin D, calcium, and bone turnover (Figure 6): First, microbial metabolites like SCFAs may promote osteoblast activity, whereas low-grade inflammation induced by gut microbial metabolites may favor osteoclast activity, thereby affecting bone remodeling, which is supported by current literature (36, 41, 42). This could explain the associations between high abundance of UCG-002 and higher LS-BMAD at baseline, high Shannon diversity and higher LS-BMAD at baseline, high abundance of *Streptococcus* and lower TBLH-BMD at baseline, as well as the low LS-BMAD after intervention, when baseline abundance of *Enterococcus* is high. In this scenario, the increased calcium demand during bone accrual could transiently lower serum calcium levels, which are rapidly normalized by homeostatic mechanisms, including PTH release, which in turn stimulates 1α-hydroxylation of 25(OH)D. Theoretically, this would also explain the high PTH and high 1,25(OH)2D plasma concentrations at baseline, when abundance of *Subdoligranulum* is high, as well as the sustained PTH after intervention, when baseline abundance of UCG-009 is high. However, the lack of observed associations between baseline *Subdoligranulum* and BMD, or between baseline UCG-009 and change in BMD during intervention, argues against this explanation.

**Figure 6:**
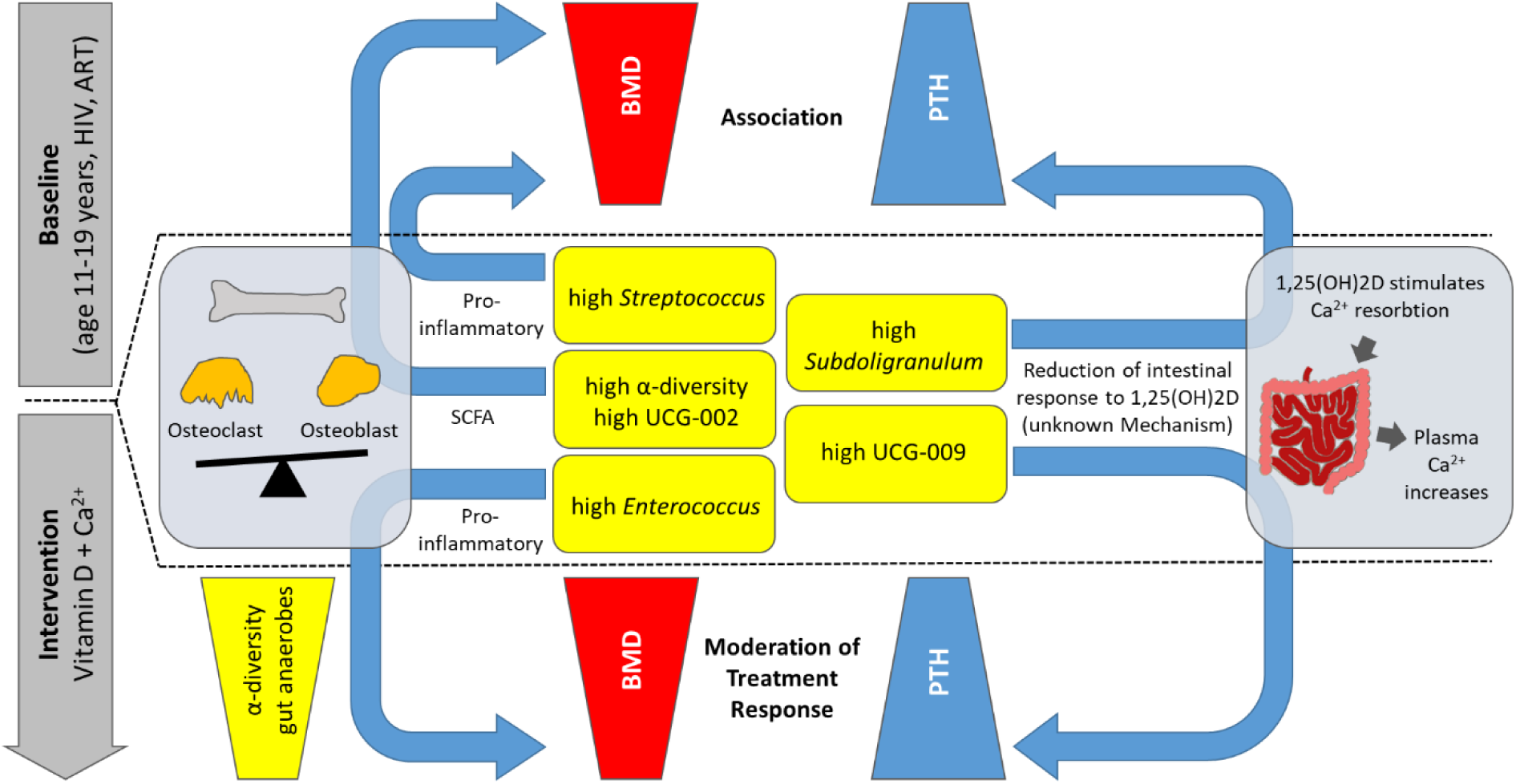
Hypothetical mechanisms linking microbiota with the regulatory network of vitamin D, calcium, and bone turnover. Two hypothetical mechanisms are displayed which could explain baseline associations and the moderation of the treatment response observed in this study. Left: Microbial metabolites like SCFAs may promote osteoblast activity, whereas low-grade inflammation induced by gut microbial metabolites may favor osteoclast activity, thereby affecting bone mineral density. Right: Microbial metabolites could reduce intestinal responsiveness to 1,25(OH)2D via a yet unknown mechanism, lowering intestinal calcium absorption and triggering compensatory PTH secretion. **Abbreviations:** BMD, bone mineral density; PTH, parathyroid hormone; ART, antiretroviral therapy; SCFA, short chain fatty acids; 1,25(OH)2D, 1,25-dihydroxyvitamin D

Secondly, microbial metabolites could reduce intestinal responsiveness to 1,25(OH)2D via a yet unknown mechanism, lowering intestinal calcium absorption and triggering compensatory PTH secretion. This could explain high PTH and high 1,25(OH)2D plasma concentrations at baseline, when abundance of *Subdoligranulum* is high. Additionally, it could explain sustained PTH after intervention, when baseline abundance of UCG-009 is high.

Microbial metabolites could also affect vitamin D uptake or 1α-hydroxylation of 25(OH)D, which is supported by previous reports.(43, 44) While this would explain the observed positive baseline associations of *Subdoligranulum* with 1,25(OH)2D concentrations, the concomitant positive association with PTH argues against this explanation.

Limitations of the study are the use of rectal swabs instead of stool samples, as well as 16S rRNA-instead of metagenomic sequencing, which limits the taxonomic resolution. Season was almost completely collinear with study site, therefore mitigating the possibility to investigate seasonal effects. Further, the lack of microbiome data from a comparable cohort without HIV precluded investigations about the role of HIV in microbiota-mediated reduced vitamin D concentrations and bone accrual in adolescents.

In summary, the microbiota-focused sub-study of the VITALITY clinical trial revealed associations between the microbiota and changes in vitamin D metabolism, PTH, and bone mineral density at baseline. Intervention by vitamin D + calcium supplementation caused a reduction of a member of the Clostridia class and, by trend, a reduction of further Clostridia members. This reduction occurred alongside an increased susceptibility to respiratory tract infections, potentially due to lowered SCFA production causing downstream immunomodulatory effects. Finally, the microbiota moderated the vitamin D + calcium supplementation mediated changes in LS-BMAD in the treatment arm of the entire cohort, and the change in PTH in participants with 25(OH)D insufficiency at baseline. In conclusion, vitamin D and calcium supplementation in adolescents living with HIV influence the composition of the gut microbiota, while, conversely, the microbiota moderate the effects of supplementation.

## Data Availability

The datasets generated during the current study will be made publicly available in the NCBI repository following peer review and publication.

## Author Contributions

RF, UES, LK, VS, CLG, SF, KK, and SLRJ conceptualized the VITALITY trial. UES and MH conceptualized the microbiota-directed sub study. MH, TM, GT, MJT, MMC, CL, and KS were responsible for data acquisition and laboratory analyses. Supervision was provided by UES, RF, PK, NVD, and MH. VS curated and validated the primary study data. MH performed the formal analyses with help from VS. Project administration was managed by RF, LK, and KK. MH wrote the original draft of the manuscript, and all authors contributed to reviewing and editing the final version.

## Use of AI tools

The authors used OpenAI’s ChatGPT (GPT-4o) to assist with R programming syntax (e.g., code structuring and debugging), without influencing the choice of analyses or statistical models. ChatGPT was also used to support English phrasing during manuscript preparation. All scientific content, interpretations, and conclusions were developed independently by the authors.

## Acknowledgements

The authors would like to thank Kristine Hagens for performing the 16S rRNA gene sequencing.

---

**Figure S1:**
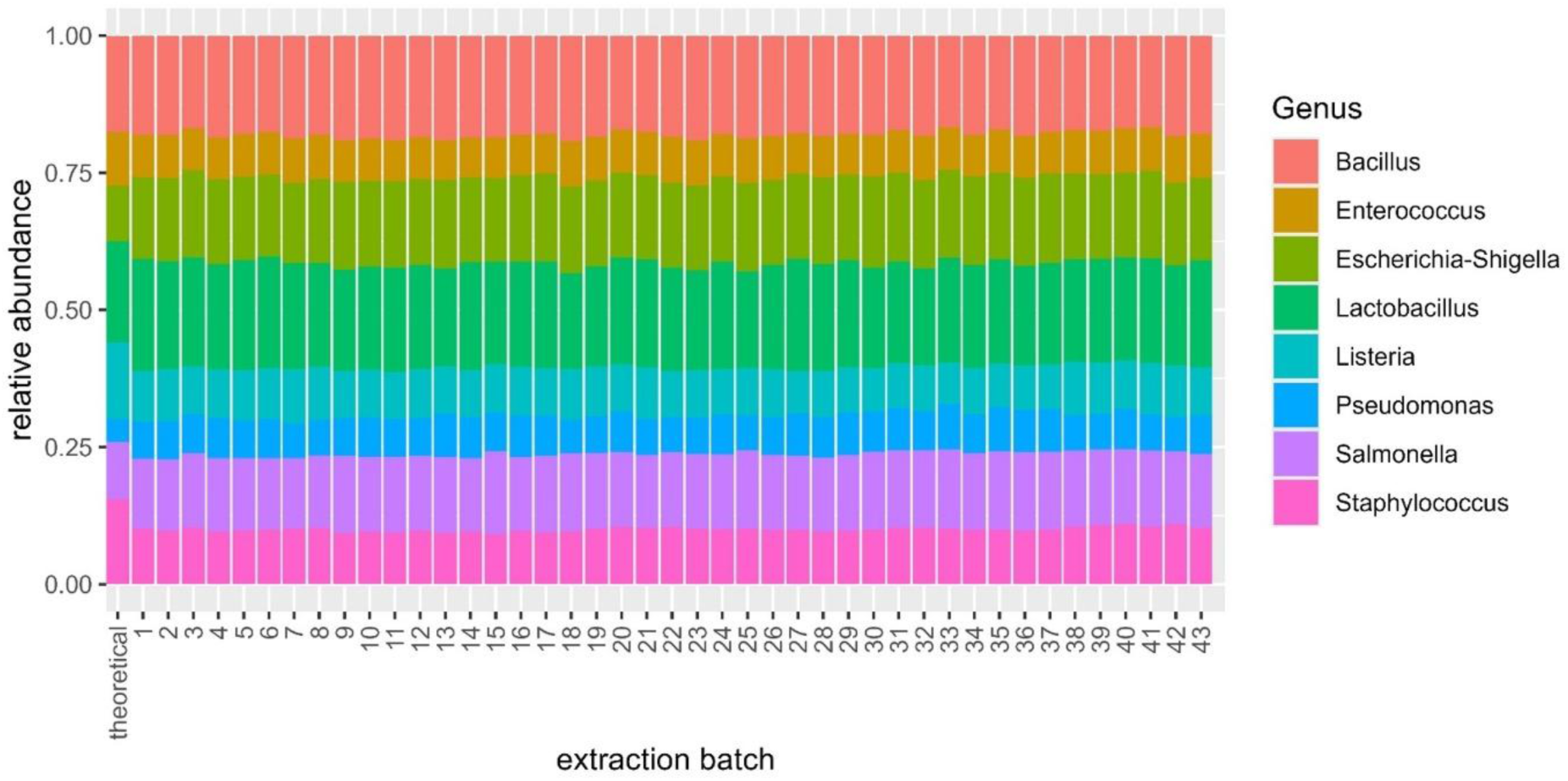
16S sequencing data of mock communities. The relative abundances of the indicated genera in mock control samples are shown across extraction batches alongside the theoretical composition.

**Figure S2:**
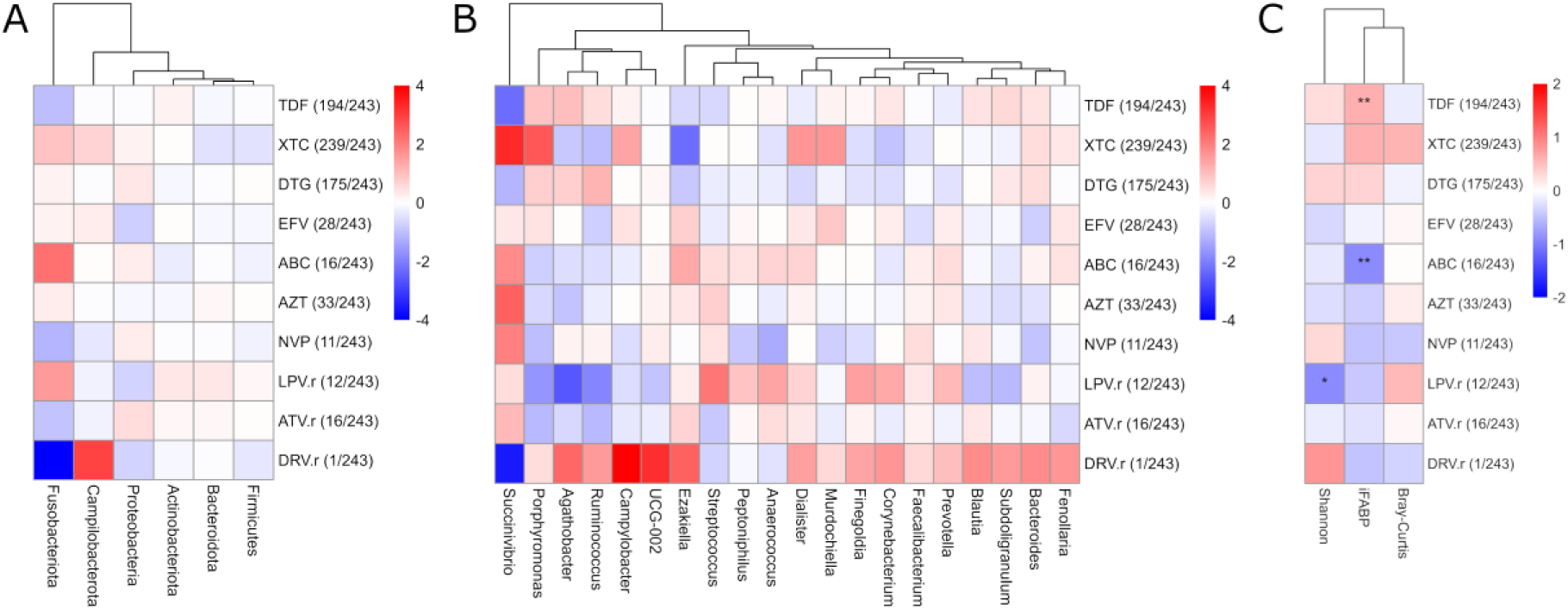
Associations of microbiota phyla and genera with the different HIV drugs prescribed. **A, B)** Differential abundances in baseline microbiota by prescribed antiretroviral drugs were assessed using LinDA, adjusting for study site. Results are shown for the 6 most abundant phyla **(A)** and the 20 most abundant genera **(B)**. Colors represent log₂ fold changes. **C)** Associations of values for Shannon diversity, Bray-Curtis dissimilarity, and plasma iFABP with prescribed antiretroviral drugs were assessed using linear regression, adjusting for study site. Colors represent standardized β-coefficients. **A-C)** The number of participants taking each drug and the total number of participants are shown in brackets. One participant with unknown therapy regimen was excluded from all analyses. **Sample size**: n=243. Asterisks indicate FDR-adjusted significance concentrations: * q<0.05, ** q<0.01 **Abbreviations**: LinDA, Linear Models for Differential Abundance Analysis; TDF, Tenofovir Disoproxil Fumarate; XTC, Emtricitabine or Lamivudine; DTG, Dolutegravir; EFV, Efavirenz; ABC, Abacavir; AZT, Zidovudine; NVP, Nevirapine; LPV.r, Lopinavir/Ritonavir; ATV.r, Atazanavir/Ritonavir; DRV.r, Darunavir/Ritonavir; iFABP, intestinal fatty acid binding protein.

**Figure S3:**
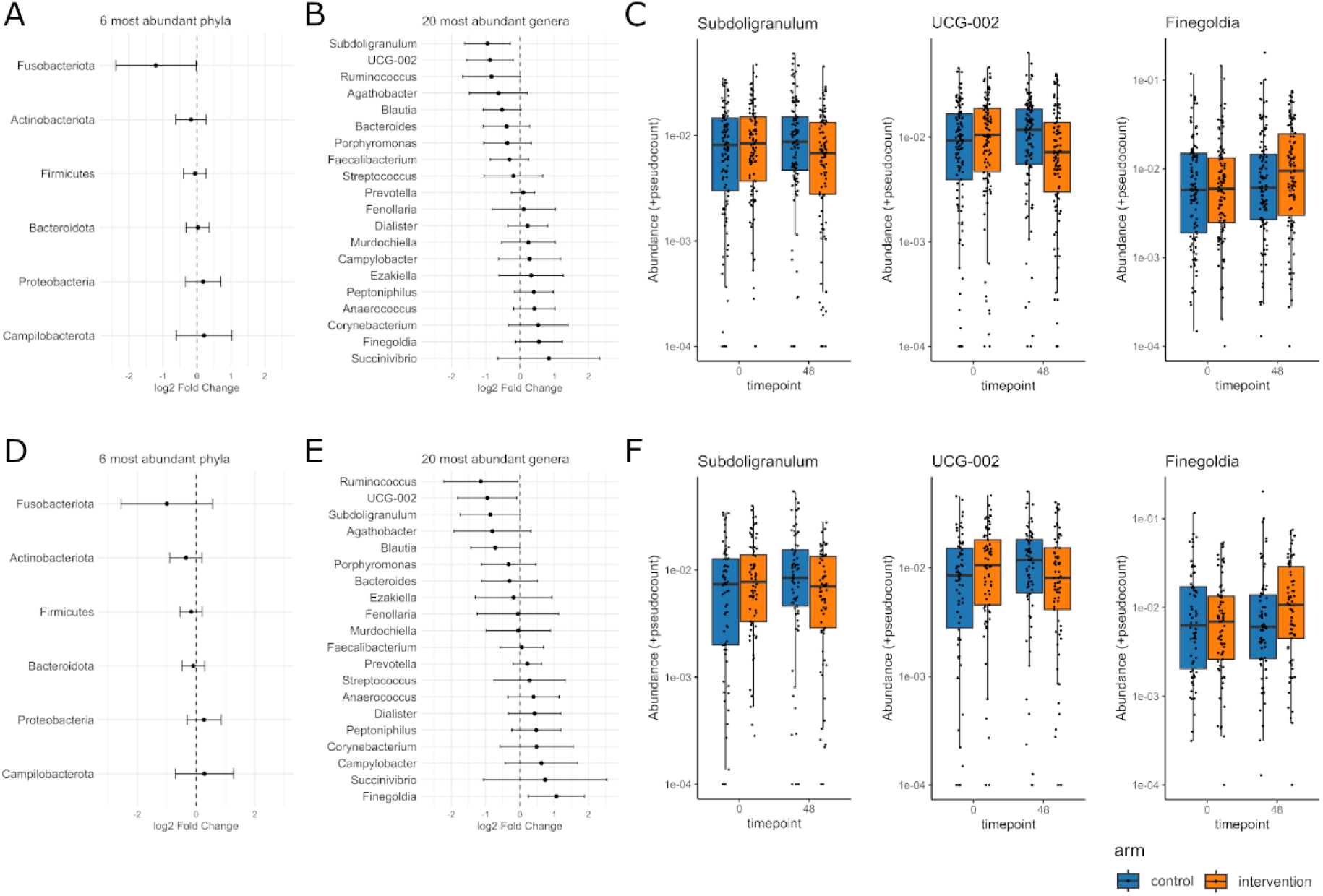
Effects of vitamin D + calcium supplementation on microbiota genera. Analyses include all participants with baseline and 48-week samples **(A-C)** or participants with 25(OH)D-insufficiency at baseline **(D-F)**. **A, B, D, E)** Differential genus-level abundances in rectal swab microbiota, associated with treatment, were assessed using LinDA, based on treatment by arm interaction, adjusted for time point, arm, and study site as fixed effects and participant as a random effect to account for repeated measures. Caterpillar plots display log₂ fold changes and 95% confidence intervals for the six most abundant phyla (**A, D**) or the 20 most abundant genera (**B, E**). The vertical line indicates no change (log₂ fold change=0). No significant associations were detected (all q > 0.05). **C, F)** Boxplots illustrate relative abundances of indicated genera across time points and treatment arms. **Sample sizes: A-C)** n=438 samples (219 participants); **D-F)** n=286 samples (143 participants) **Abbreviations:** LinDA, Linear Models for Differential Abundance Analysis.

**Figure S4:**
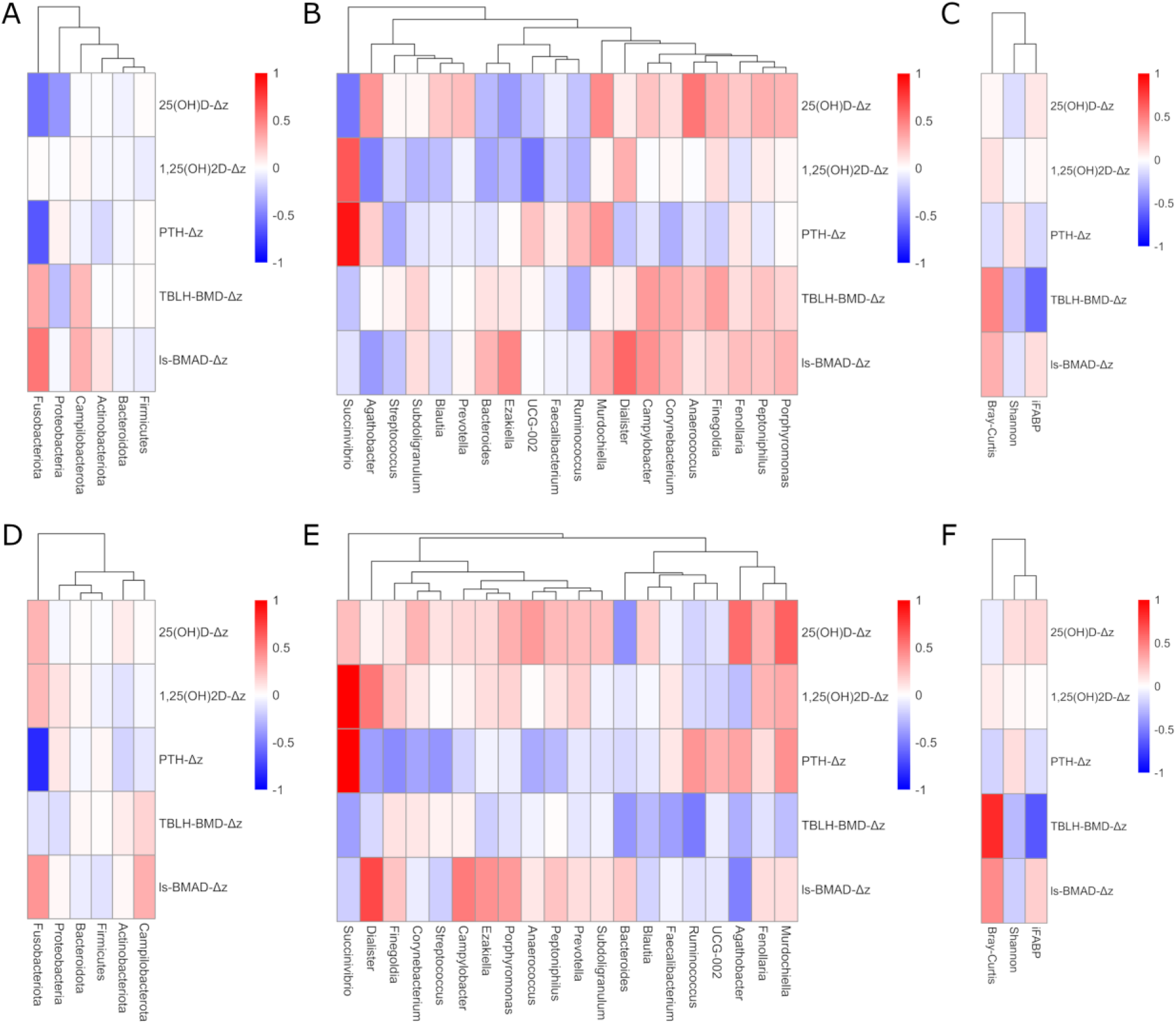
Predictors of treatment response to vitamin D + calcium supplementation. Analyses include all participants in the intervention arm **(A-C)** or those in the intervention arm with 25(OH)D-insufficiency at baseline **(D-F)**. Variables 25(OH)D, 1,25(OH)2D, PTH, TBLH-BMD Z-score, and LS-BMAD Z-score were Z-transformed and changes in Z-scores between 48-weeks and baseline samples (ΔZ) were used as proxies for treatment response. **A, B, D, E)** Associations between genus-level abundance at baseline and ΔZ of the indicated variables were assessed using LinDA, adjusted for study site. Heatmaps indicate log_2_ fold changes for the 6 most abundant phyla **(A, D)** or the 20 most abundant genera **(B, E)**. **C, F)** Associations between Shannon diversity, Bray-Curtis dissimilarity, and plasma iFABP and ΔZ of the indicated variables were assessed using linear regression models, with study site included as a covariate. Heatmaps indicate standardized β-coefficients. **Sample sizes**: A-C: n=107-109 and D-F: n=73-74 participants due to incomplete TBLH-BMD Z-score and LS-BMAD Z-score data. **Abbreviations**: LinDA, Linear Models for Differential Abundance Analysis; iFABP, intestinal fatty acid binding protein; PTH, parathyroid hormone; TBLH-BMD-Z, total body less head bone mineral density Z-score; LS-BMAD-Z, lumbar spine bone mineral apparent density Z-score.

